# Oligodendrocyte precursor cells guide the migration of cortical interneurons by unidirectional contact repulsion

**DOI:** 10.1101/2021.05.27.446000

**Authors:** Fanny Lepiemme, Gabriel Mazzucchelli, Carla G. Silva, Laurent Nguyen

**Author notes:** Correspondence: Laurent Nguyen,; Carla Silva. Equal contribution (Carla G. Silva, Laurent Nguyen).

## Abstract

The cerebral cortex is built by neural cells that migrate away from their birthplace. In the forebrain, ventrally-derived oligodendrocyte precursor cells (vOPCs) travel tangentially together with cortical interneurons (cINs) to reach the cortex. After birth, vOPCs form transient synapses with cINs before engaging later into myelination. Here we tested whether these populations interact during embryogenesis while migrating. By coupling histological analysis of mouse genetic models with live imaging, we showed that, while responding to the chemokine Cxcl12, vOPCs and cINs occupy mutually-exclusive forebrain territories. Moreover, vOPCs depletion selectively disrupts the migration and distribution of cINs. At the cellular level, we found that by promoting unidirectional contact-repulsion (UCoRe) of cINs, vOPCs steer their migration away from blood vessels and contribute to their allocation to proper migratory streams. UCoRe is thus an efficient strategy to spatially control the competition for a shared chemoattractant, thereby allowing cINs to reach proper cortical territories.

## Introduction

Oligodendrocytes differentiate from oligodendrocyte progenitor cells (OPCs) and populate the entire CNS. After birth, they progressively wrap axons to support rapid saltatory conduction of action potentials as well as to provide metabolic support to neurons via axo-myelinic channels ^1^. Some OPCs convert into Ngn2 glia, which is the most proliferative pool of cells in the mature brain ^2^. Importantly, these cells also form bona fide synapses with neurons ^3–5^. In the forebrain, OPCs have multiple origins and are generated in successive “spatio-temporal waves” that populate the cortex from embryonic development onward ^6^. The two first OPC waves (vOPCs) are produced in the ventral forebrain and are followed by a perinatal one born in the cerebral cortex (dOPC), which gradually expands and outnumbers the others ^6^. OPCs from different origins are genetically distinct and several intermediates coexist during development ^7^, but they further converge into similar transcriptional states after birth ^8^. While the second and third OPC-waves mainly contribute to axon myelination, the function of the first vOPC cohort remains elusive. This cell population was first thought to be transient and eliminated at birth ^6^, but recent single cell analyses showed its progressive dilution in the maturing brain and persistence in selective brain regions during adulthood ^8^. Most vOPCs share common embryonic origins with cortical interneurons (cINs). They are born from progenitors of the ganglionic eminences (GEs) and the preoptic area (POA), where genetic cross-inhibitory mechanisms control the balanced generation of cINs and vOPCs. The expression of some Dlx homeobox transcription factors drives cIN fate by repressing the OPC gene *Olig2*^9^, while Olig1 expression promotes the specification of vOPCs and prevents the acquisition of a cIN fate by preventing the expression *Dlx1/2* ^10^. vOPCs from the POA (Dbx1-positives) or the MGE (Nkx2.1-positive) clustered with their lineage-related cINs in the cortex of newborn mice where they form transient synapses, before engaging later into myelination of specific local circuits ^5, 11^. During embryogenesis, vOPCs an cINs also migrate concomitantly into the cerebral cortex ^6, 12^, a process during which these cells may interact and cooperate. Here, we tested this hypothesis by performing live imaging on organotypic forebrain slices from transgenic mice to analyze the distribution and dynamic behavior of contemporary vOPCs and cINs. We first showed that, despite coexisting in time and space, these cells tend to occupy different territories and to invade the cortex by using distinct migration modes and paths. The elimination of vOPCs from early forebrain results in cINs distribution and migration parameters changes, supporting the existence of a functional crosstalk between these two cell types from the onset of corticogenesis. At the cellular level, we found that vOPCs repulse cINs by unidirectional contact repulsion, a process that we named UCoRe. This peculiar behavior contribute to the guidance of cINs in stream and also prevent them to cluster around blood vessels (BVs). Indeed, vOPCs and cINs express Cxcr4 and are both responsive to the chemokine Cxcl12 released by BVs that promotes OPCs migration and proliferation on these structures^13^. We found that vOPC elimination frees cortical BVs and locally changes the availability of Cxcl12, thereby altering cINs migration in subpallial territories and in the cortical wall. Altogether, our study demonstrates that vOPCs help cINs to navigate across diverse substrates found along their way to the cerebral cortex by using UCoRe. This unprecedented mode of repulsion represents an efficient strategy to allow distinct cell types, which are responsive to a common chemoattractant, to progress distinctly on shared forebrain territories.

## Results

### Spatial segregation of migrating cINs and vOPCs in the ventral forebrain

We performed immunolabeling of E11.5 and E13.5 embryos to study the distribution pattern of first-wave Pdgfrα^+^ vOPCs ^6^ and early migrating cINs in the forebrain (derived from the POA and the ganglionic eminences, or GEs) that express calbindin (CB)^14^ (we did not detect any cell co-expressing both markers). At E11.5, both cell types mostly distribute into non-overlapping subpallial territories (Figures 1A–1D). vOPCs populate the germinal zones of the medial (M) GE/POA and cINs are distributed throughout the subventricular zone (SVZ) and mantle zone (MAZ). Two days later, vOPCs mostly accumulate in the SVZ and along the MAZ, against a stream of migrating cINs (Figures 1E–1H). At this stage, vOPCs are also mostly detected in the ventricular zone (VZ) and SVZ of the MGE and the POA, where they are actively generated (Figure 1F). The CB^+^cINs born in the MGE organize into a deep migratory stream (DMS), and those derived from the POA form a superficial migratory stream (SMS) (Figure 1E) ^15^. Interestingly, vOPCs do not integrate these migratory streams, suggesting that during early embryonic periods, vOPCs and cINs are spatially segregated in the subpallium (Figure 1B). This is further confirmed by the analysis of maximum fluorescence intensities (MFI) and relative number of cells expressing Pdgfra or CB at E11.5 (Figures 1C–1D) and at E13.5 (Figure 1G–1H), supporting a spatial segregation of vOPCs and cINs populations in the ventral forebrain. We showed that vOPCs that occupy the MGE/POA territory above the DMS mainly arise from Nkx2.1^+^ (87.72%% ± 5.04%) and Dbx1^+^ (31.23 % ± 2.05%) progenitors (Figures 1I–1L). Time lapse video microscopy analysis performed in this territory on organotypic slices from E13.5 mice showed that the majority of GFP^+^ vOPCs are highly dynamic with some moving away to initiate their migration towards the developing cortical wall (Supplemental video 1). These data demonstrate that contemporary vOPCs and cINs populate distinct subpallial territories during embryogenesis through the dynamic formation of boundaries.

**Figure 1.**
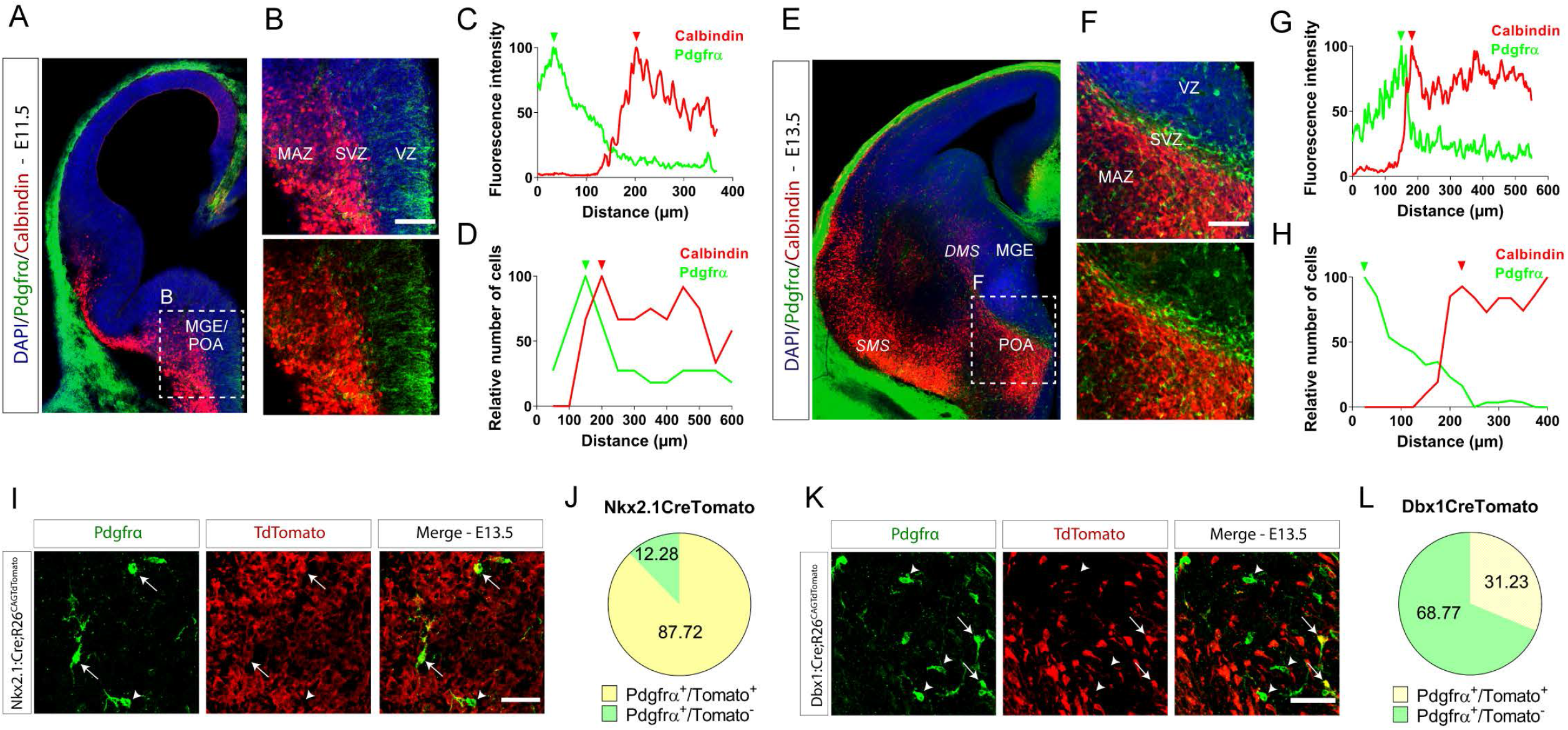
vOPCs and cINs occupy distinct subpallial territories. (A-B) Immunolabeling of vOPCs and cINs in a coronal brain slice of a WT E11.5 mouse embryo. vOPCs express Pdgfrα (green) and cINs express calbindin (red). The squared region (white dashed line) corresponds to a MGE/POA magnified area represented in (B). (B) vOPCs (green) localize mostly in VZ and cINs (red) in SVZ and MAZ. Nuclei are counterstained with DAPI (blue – upper panel). Scale bar: 100μm. (C) Fluorescence intensity distribution of Pdgfrα (green) and calbindin (red) immunolabelings within the magnified area represented in (B). Maximum fluorescence intensities are pointed by color-matching arrowheads for Pdgfrα (green) and calbindin (red) immunolabelings. (D) Distribution of relative numbers of Pdgfrα^+^ and calbindin^+^ cells observed in (B). Maximum relative numbers are pointed by color-matching arrowheads for Pdgfrα^+^ (green) or calbindin^+^ (red) cells. (E) Immunolabeling of vOPCs and cINs in a coronal brain slice of a WT E13.5 mouse embryo. vOPCs express Pdgfrα (green) and cINs express calbindin (red). (F) Magnified MGE/POA area squared in (E), where vOPCs (green) localize at the transition between the VZ and SVZ and cINs (red) in the MAZ of the MGE and POA. Nuclei are counterstained with DAPI (blue - upper panel). Scale bar: 100μm. (G) Fluorescence intensity distribution of Pdgfrα (green) and calbindin (red) immunolabelings within the magnified area represented in (F). Maximum fluorescence intensities are pointed by color-matching arrowheads for Pdgfrα (green) and calbindin (red) immunolabelings. (H) Distribution of relative numbers of Pdgfrα^+^ and calbindin^+^ cells observed in (F). Maximum relative numbers are pointed by color-matching arrowheads for Pdgfrα^+^ (green) or calbindin^+^ (red) cells. (I) Lineage tracing of Pdgfrα^+^ vOPCs in Nkx2.1:Cre;R26^CAGTdTomato^ at E13.5. White arrows point Pdgfrα^+^ cells generated by Nkx2.1^+^ progenitors. The white arrowhead points a Pdgfrα^+^ vOPC originating from a Nkx2.1^-^ progenitor. Scale bar: 50μm. (J) Proportion of Pdgfrα^+^ cells originating from Nkx2.1^+^ progenitors (yellow) or from other progenitor cells (green). (K) Lineage tracing of Pdgfrα^+^ vOPCs in Dbx1:Cre;R26^CAGTdTomato^ at E13.5. White arrows point Pdgfrα^+^ cells generated from Dbx1^+^ progenitors. White arrowheads point Pdgfrα^+^ cells originated from Dbx1^-^ progenitor. Scale bar: 50μm. (L) Proportion of Pdgfrα^+^ cells originating from Dbx1^+^ progenitors (yellow) or from other progenitor cells (green).

### cINs and vOPCs use different migration strategies to invade the developing cortical wall

We performed immunolabeling to study the distribution of vOPCs and cINs in the developing cortical wall of E13.5 mice and we followed their migration in transgenic mice (Dlx5,6:Cre-GFP for cINs and Sox10:loxGFP-STOPlox-DTA - further refereed as Sox10:GFP-DTA - for vOPCs). While cINs migrate within two main tangential streams - one in the marginal zone (MZ) and the other in the intermediate zone (IZ) -, vOPCs do not display stream organization (Figure 2A; Supplemental videos 2-3). At 16.5, numerous cINs undergo intracortical dispersion and invade the cortical plate (CP), while most vOPCs accumulate under the CP, where thalamocortical fibres extend (Figures 2B, 2C). However, at this developmental stage, some vOPCs have already invaded the CP by climbing along BVs, as previously reported, following a ventral to dorsal gradient (Figure 2D) ^13^. Although cINs do not preferentially migrate on BVs, they can be detected in their vicinity ^16^. We observed that the percentage of vOPCs migrating on BVs increases as corticogenesis proceeds, following the temporal progression of cortical angiogenesis ^17^ (E13.5: 52.46% ± 4.45%, E16.5: 77.52%±7.86%; p=0.032; Figures 2E, 2F). vOPCs use comparable strategies to move in the parenchyma or along BVs, suggesting that their migration mode mostly relies on intrinsic mechanisms (Figures S1A–S1D).

**Figure 2.**
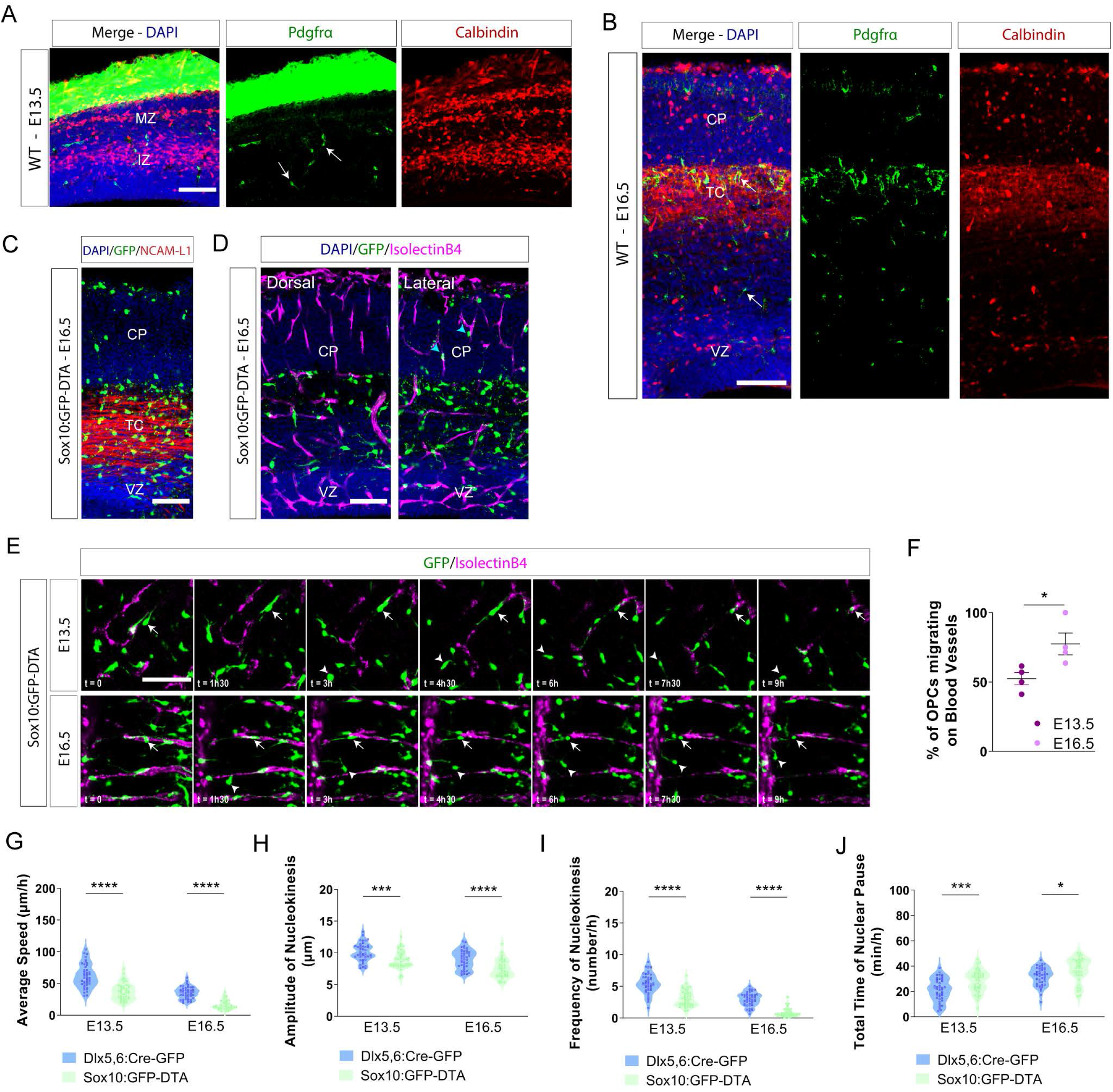
Migrating vOPCs and cINs display distinct distributions in pallial territories. (A) Distribution of vOPCs (Pdgfrα^+^, green) and cINs (calbindin^+^, red) in the cortical wall of a WT E13.5 embryo (25μm Z-stack). cINs distribute within two migratory streams, located in the MZ and IZ. The white arrows in the middle panel point vOPCs, distributed randomly in cortical territories. The left panel is the merge between the middle and right panels. Nuclei are counterstained with DAPI. The intense green staining in extracortical territories corresponds to expression of Pdgfrα in meningeal cells. Scale bar: 100μm. (B) Relative distribution of Pdgfrα^+^ vOPCs (green) and calbindin^+^ cINs (red) in the cortical wall of a WT E16.5 embryo. The cortical plate (CP) is mainly populated by calbindin^+^ cINs, while Pdgfrα^+^ vOPCs cells accumulate underneath the CP and above the VZ. The left panel is the merge between the middle and right panels. Nuclei are counterstained with DAPI. Scale bar: 100μm. (C) Immunolabeling of a brain section of a E16.5 Sox10:GFP-DTA embryo showing the accumulation of vOPCs (green) on NCAM-L1^+^ thalamocortical fibers (TC; red). Nuclei are counterstained with DAPI. Scale bar: 100μm. (D) Dorsal (left) and lateral (right) sections of an E16.5 Sox10:GFP-DTA brain showing the density of vOPCs in the TC region between the CP and the VZ. vOPCs invasion of the CP via BVs (purple staining) starts in lateral section but not in dorsal section. Scale bar: 100μm. (E) Timeline of 9h showing vOPCs migrating on BVs (white arrows) or within brain parenchyma (white arrowheads) in cortical regions of a Sox10:GFP-DTA at E13.5 and E16.5. Scale bar: 50μm. (F) Fraction of vOPCs migrating on BVs in the cortical wall of E13.5 and E16.5 Sox10:GFP-DTA embryos. The fraction of vOPCs migrating on BVs increases at E16.5 (n = 61-70 cells in at least 3 embryos from 3 females; unpaired t-test, *p=0.032). (G-J) Violin plots representing the migration parameters of Dlx5,6:Cre-GFP interneurons and Sox10:GFP-DTA vOPCs in organotypic brain slices at E13.5 and E16.5. The average speed of migration (G), amplitude of nucleokinesis (H) and frequency of nucleokinesis (I) are higher in cINs as compared to vOPCs both at E13.5 and E16.5 (n = 38-44 cells from at least 3 embryos from 3 dams; two-way ANOVA; ****p<0.0001, ***p=0.0008 and ****p<0.0001 at E13.5; ****p<0.0001, ****p<0.0001 and ****p<0.0001 at E16.5). The total time of nuclear pause (J) is longer in vOPCs at both stages (n = 38-44 cells in at least 3 embryos from 3 females; two-way ANOVA, ***p=0.0008 at E13.5 and *p=0.038 at E16.5).

At the cellular level, both cINs and vOPCs undergo saltatory migration but vOPCs move at slower speed, as a result of lower nucleokinesis frequency and amplitude (Figures 2G–2I) combined with longer pausing times (Figure 2J). Altogether, these results show that contemporary vOPCs and cINs use different migration modes and substrate to invade the cortical wall.

### The genetic elimination of vOPCs alters the distribution pattern of cINs in the forebrain

First-wave vOPCs have been reported to display preferential interaction with lineage-related cINs in order to form synaptically-coupled vOPCs-cINs clusters at birth, which later contribute to cortical myelination ^5, 11^. Here, we wondered if vOPCs and cINs initiate some interaction during embryogenesis and whether this would be important for their distribution and migration within the cerebral cortex. To test this hypothesis, we crossed Pdgfτα-Cre^ERT^ and Sox10:GFP-DTA mice to generate embryos where vOPCs are totally eliminated upon intraperitoneal injection of tamoxifen (tam ip; 100% of depletion, n=13) in pregnant dams (OPC-depleted embryos) (Figure 3A). Successive tam ip at E11.5 and E12.5 eliminated first-wave vOPCs by E13.5, without inducing bystander cell death, change in GE progenitor proliferation rate, or in cINs numbers (Figures S1E–S1K). The analysis of the forebrain of E14.5 OPC-depleted embryos revealed that, conversely to the SMS, the DMS enlarged as a result of an increased dispersion of CB^+^ cINs (Figures 1E, 3B). These cells invade the MGE SVZ, therefore leading to a reduced distance between the MGE ventricle limit and the internal DMS border (Figures 3B, 3C; controls: 230.30μm ± 6.76μm; OPC-depleted: 132.50μm ± 7.58μm, p=0.029). Analysis of the dorsal forebrain of E13.5 and E16.5 embryos, showed a reduced number of cINs into the cortical streams upon depletion of vOPCs (Figures 3D–3F; E13.5 rostral: control is 70.23 ± 14.13 and OPC-depleted is 55.57 ± 10.25, p=0.0024; E13.5 medial: control is 74.31 ± 15.51 and OPC-depleted is 57.29 ± 10.01, p=0.0004; E13.5, caudal: control is 75.85 ± 16.23 and OPC-depleted is 58.43 ± 10.46, p=0.0003; E16.5 rostral: control is 101.80 ± 8.54 and OPC-depleted is 62.71 ± 9.96, p<0.0001; E16.5 medial: control is 92.00 ± 20.83 and OPC-depleted is 58.00 ± 13.85, p=0.0007; E16.5, caudal: control is 98.33 ± 5.89 and OPC-depleted is 61.43 ± 13.55, p<0.0001). Despite the progressive increase of vOPC numbers generated by the second wave in OPC-depleted cortex (successive tam ip at E11.5 and E12.5 to eliminate first-wave vOPCs), the number of cINs in the cortical wall remains low around birth, suggesting a long-lasting phenotype (GFP^+^cINs-control: 187.5 ± 17.95 and GFP^+^cINs-OPC-depleted: 99.5 ± 17.13; p=0.0008; Figures S1L–S1M). The migration defect impacts most cINs (GFP^+^cINs-control: 40.0 ± 3.34 and GFP^+^cINs-OPC-depleted: 22.25 ± 2.72; p=0.029; Figures S1N–S1O) and is specific since we did not observe any cortical distribution change of microglia (Figures S1P–S1Q), which itself control the final distribution of cINs ^18^. Moreover, depleting cINs by expressing DTA in Dlx5.6^+^ cells had no effect on the number or localization of vOPCs within the cortical parenchyma (Figure S1R–S1S). Altogether, these results suggest that first-wave vOPCs influence selectively the forebrain distribution and sorting of cINs in the cerebral cortex.

**Figure 3.**
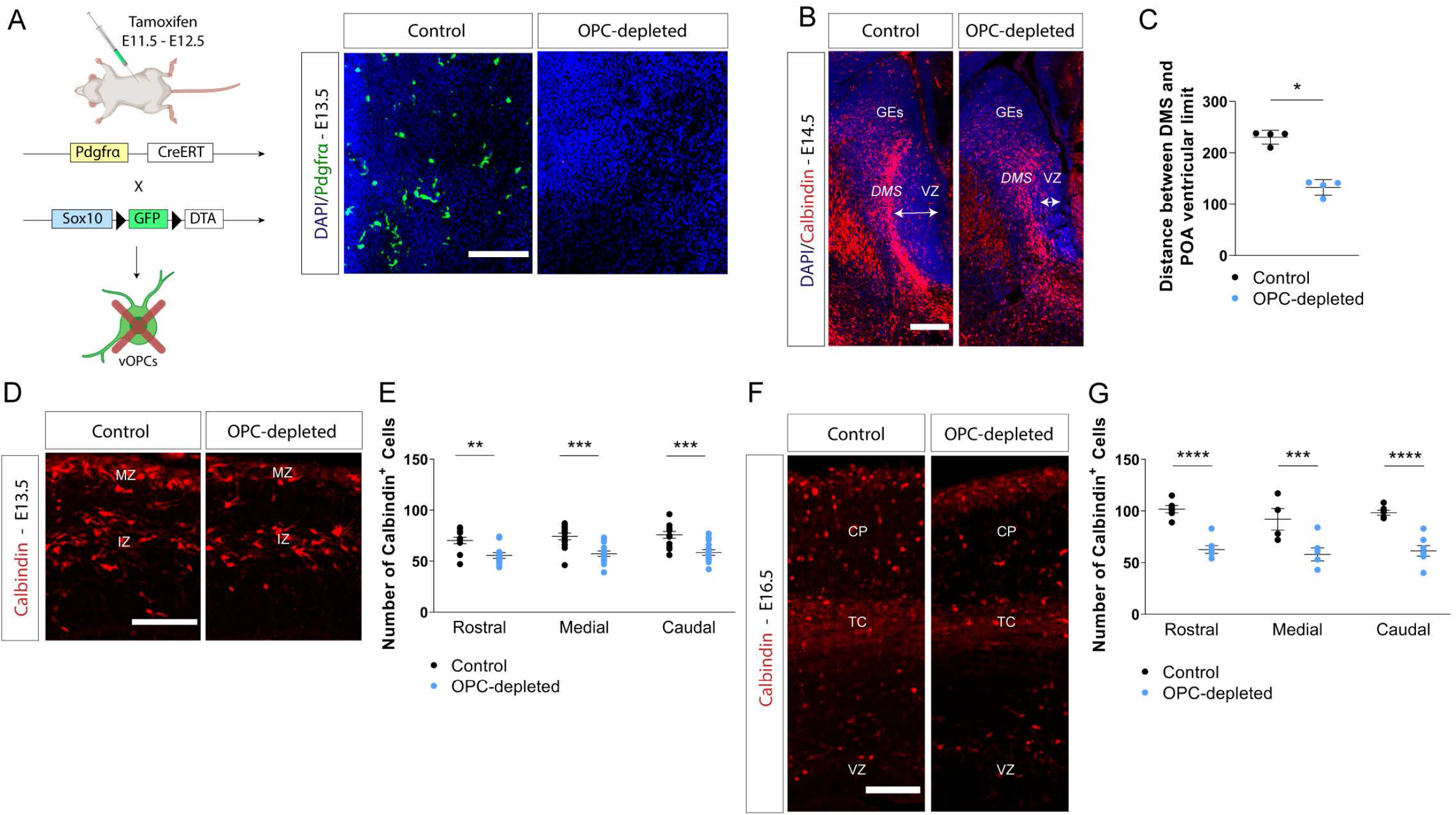
First-wave vOPCs depletion alters cINs distribution in pallial and subpallial territories. (A) Mouse breeding strategy to conditionally express subunit A of diphteria toxin (DTA) and eliminate Pdgfrα^+^:Sox10^+^ first-wave vOPCs upon tamoxifen administration. Immunolabeling of brain slices from E13.5 Sox10:GFP-DTA embryo (left) or OPC-depleted (right; Pdgfrα:Cre^ERT^;Sox10:GFP-DTA) embryo shows that tamoxifen administration at E11.5 and E12.5 leads to a complete elimination of first-wave vOPCs (green; n= 13 embryos from 6 females, 100% of depletion). Nuclei are counterstained with DAPI. Scale bar: 50μm. (B) Immunolabeling of calbindin^+^ cINs (red) and Pdgfrα^+^ vOPCs (green) in the subpallium of E13.5 Control (Pdgfrα:Cre^ERT^) and OPC-depleted embryos. Migrating calbindin^+^ cINs organized into a DMS stream, which enlarged in OPC-depleted embryo. Scale bar: 200μm. (C) Quantification of the distance between the DMS internal border filled with calbindin^+^ cINs (red) and the POA ventricular limit, as shown in (B) (n = 4 embryos *per* group from 3 females; Mann-Whitney test, *p=0.029). (D) Immunolabeling of calbindin^+^ cINs (red) in the cortical wall of E13.5 Control (left) and OPC-depleted (right) embryos. The number of migrating cINs in the MZ and IZ streams is decreased upon vOPCs elimination. Scale bar: 50μm. (E) Quantification of the number of calbindin^+^ cINs (red) in rostral, medial and caudal regions of the cortex from Control (black dots) and OPC-depleted (blue dots) embryos (n= 13-14 embryos from 6 females; two-way ANOVA, **p=0.0024, ***p=0.0004, ***p=0.0003). (F) Immunolabeling of calbindin^+^ cINs (red) in the cortical wall of E16.5 Control (left) and OPC-depleted (right) embryos. The number of calbindin^+^ cINs (red) in the cortical plate (CP) is decreased. Scale bar: 100μm. (G) Quantification of the number of calbindin^+^ cINs (red) in rostral, medial and caudal regions of the cortex of E16.5 Control (black dots) and OPC-depleted (blue dots) embryos (n = 6-7 embryos from 4 females; two-way ANOVA, ****p<0.0001, ***p=0.0007, ****p<0.0001).

### Loss of vOPCs affects cINs migration in the cortical wall

Here, we tested whether the impaired forebrain distribution of cINs upon vOPCs loss results from a change in their migration behavior. To test this hypothesis, we crossed OPC-depleted with Dlx5,6:Cre-GFP mices to generate embryos (GFP^+^cINs-OPC-depleted) to follow the migration of GFP^+^ cINs (Figure 4A). We performed live imaging of organotypic brain slices from E13.5 embryos and showed a reduction of cortical invasion by GFP^+^ cINs in GFP^+^cINs-OPC-depleted embryos, as compared to their controls (Figures 4B–4C). These cINs were migrating slower (GFP^+^cINs-OPC-depleted: 17.85 μm/h ± 2.18 μm/h and GFP^+^cINs-control: 50.68 μm/h ± 2.90 μm/h; p≤0.0001; Figure 4D) and closer to isolectinB4-labelled BVs, as compared to their controls (travelled distance 3h after first contact with BVs; GFP^+^cINs-OPC-depleted: 38.00μm ± 8.93μm and GFP^+^cINs-control: 112.70μm ± 12.63μm, p<0.0001; Figures 4B–4C, 4E). At this developmental stage, the migration parameters of cINs in cortical area devoid of BVs was comparable between GFP^+^cINs-OPC-depleted and control cortices (Figures S2A–S2D), suggesting that change in cINs migration between genotypes results from a lack of BVs coverage by vOPCs. BVs are indeed a preferential migration substrate for vOPCs in the cortex ^13^, which remained morphologically unaffected (Figures S2E–S2H). Moreover, the proteomic analysis on the cortical wall of E13.5 OPC-depleted embryos and their control did not reveal molecular changes associated with BVs (Figures S2I–S2J). cIN migration into the cortical wall requires gradients of chemoattractant molecules, such as Nrg1, Cxcl12, and VEGF ^16, 19, 20^. Endothelial cells act as a discrete source of Cxcl12 ^13^, which attracts vOPCs expressing the chemokine receptor 4 (Cxcr4) upon activation of Wnt signalling ^13^. By combining proteomic and ELISA analyses, we did not detect changes of expression and release of Cxcl12 in cortices from E13.5 OPC-depleted and controls embryos (Figures S2J–S2K). We thus postulated that cINs, which also express Cxcr4^21^, would migrate closer to vessels as a result of an increase availability of endothelial Cxcl12 in the absence of covering vOPCs. To test this hypothesis, we incubated organotypic brain slices from E13.5 GFP^+^cINs-OPC-depleted and their control embryos with Cxcl12 blocking antibodies (AbCxcl12) and we measured the distance travelled by cINs after having contacted a BV. We found that cINs spent less time interacting with BVs and migrated farther from these structures upon initial contact in GFP^+^cINs-OPC-depleted brain slices incubated with AbCxcl12 (travelled distance 3h after first contact with BVs; vehicle: 14.82 μm ± 5.71 μm and AbCxcl12: 96.31 μm ± 14.62 μm, p≤0.0001; Figures 4F). These changes correlated with an increased average speed of cINs migration in GFP^+^cINs-OPC-depleted brain slices incubated with AbCxcl12 (vehicle: 12.53 μm/h ± 2.26 μm/h and AbCxcl12: 35.01 μm/h ± 4.72 μm/h; p≤ 0.0001; Figure 4G). Incubation of control brain slices with AbCxcl12 did not modify the distance travelled by cINs from initial BVs contact (vehicle: 138.30 μm ± 10.37 μm and AbCxcl12: 139.30 μm ± 10.18 μm; p>0.99; Figure 4F). Average speed was also similar between treatment conditions (vehicle: 51.42 μm/h ± 3.45 μm/h and AbCxcl12: 48.91 μm/h ± 2.57 μm/h; p=0.95; Figure 4G). Altogether, these results suggest that the coverage of BVs by vOPCs prevents Cxcl12-mediated cINs-BVs interaction. To test this hypothesis, we induced vOPC detachment from the vasculature by blocking their Wnt signalling transduction with XAV939 in organotypic brain slices^13^. Upon detachment of vOPCs from BVs, we observed more cINs around vessels. cINs travelled reduced distance in control brain slices incubated with XAV939 (travelled distance 3h after first contact with BVs, vehicle: 127.50 μm ± 6.14 μm and XAV939: 34.29 μm ± 7.03 μm, p≤0.0001; Figure 4H), and migrate slower (average migration speed, vehicle: 44.63 μm/h ± 1.85 μm/h and XAV939: 15.22 μm/h ± 2.42 μm/h, p≤0.0001; Figure 4I). Altogether, these results suggest that the migration of vOPCs along BVs limits the interaction of cINs with the vasculature and supports a possible competition between cINs and vOPCs for the local pool of endothelial Cxcl12.

**Figure 4.**
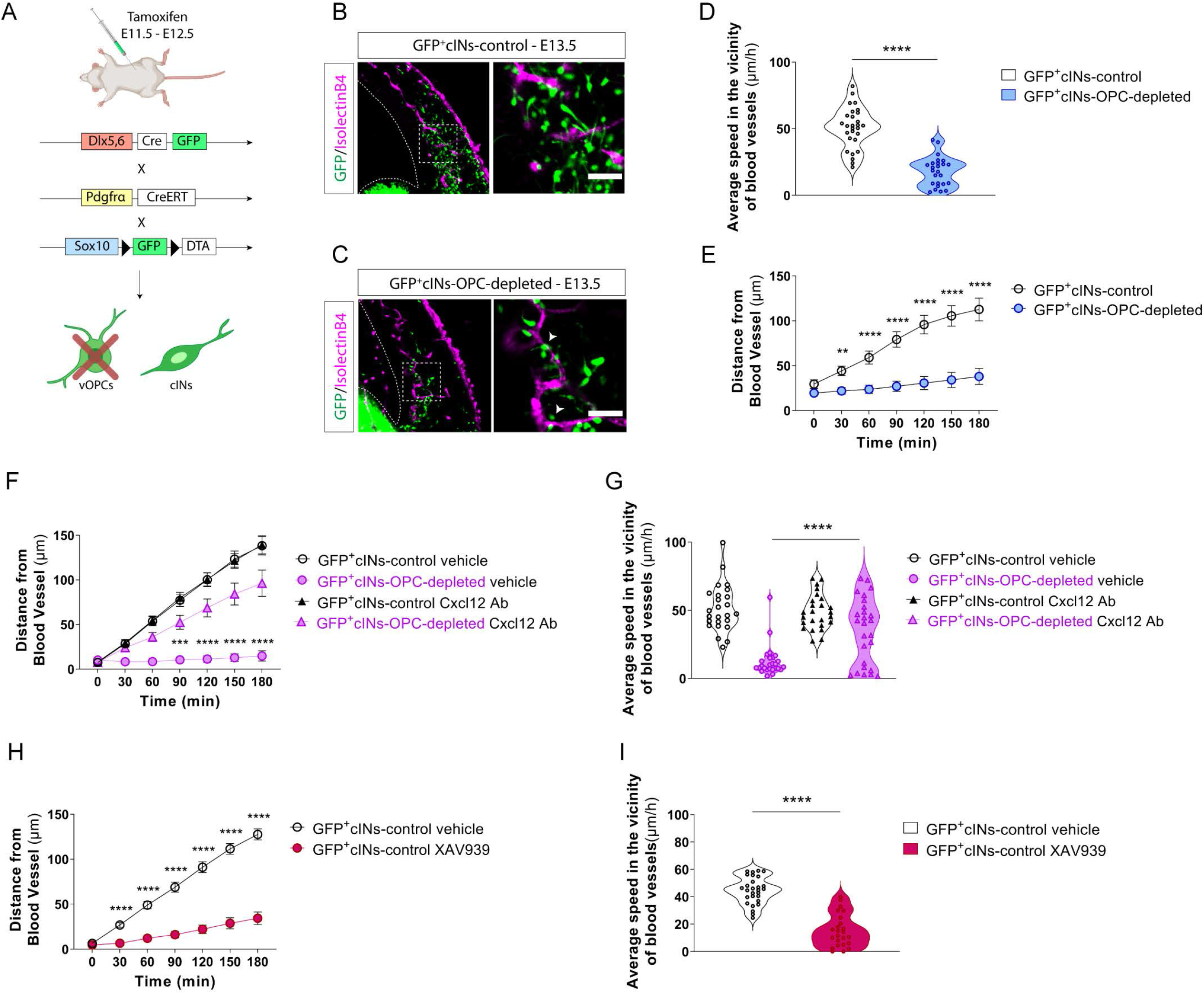
First-wave vOPCs depletion leads to changes in cIN migration behavior in vicinity to BVs. (A) Mouse breeding strategy to conditionally express subunit A of diphteria toxin (DTA) and eliminate Pdgfrα^+^;Sox10^+^ first-wave vOPCs upon tamoxifen administration and to label Dlx5,6-GFP^+^ cINs. (B-C) Time-lapse video microscopy sequence showing the distribution of Dlx5,6-GFP^+^ cINs (green) around BVs (purple) on organotypic brain slices from E13.5 GFP^+^cINs-control (B) and GFP^+^cINs-OPC-depleted embryos (C). BVs were labeled with isolectinB4 (purple) and the white arrowheads in (C) point Dlx5,6-GFP^+^ cINs located in the vicinity of BVs. Scale bar: 50μm. (D) Distribution of average migration speeds of Dlx5,6-GFP^+^ cINs moving along cortical BVs in E13.5 GFP^+^cINs-control (open circles) and GFP^+^cINs-OPC-depleted brain slices (blue circles) (n = 25-28 cells from 3 embryos, from at least 3 dams; Unpaired t-test; ****p<0.0001). (E) Histogram representing distance travelled by Dlx5,6-GFP^+^ cINs from initial contact to cortical BVs in brain slices from E13.5 GFP^+^cINs-control (open circles) or GFP^+^cINs-OPC-depleted embryos (blue circles). Time 0 corresponds to initial contact of cIN with a BV (n = 23 cells from 3 embryos from 3 dams; two-way ANOVA; **p=0.0048 after 30 min, ****p<0.0001 after 60 to 180 min). (F) Histogram representing the distance travelled by Dlx5,6-GFP^+^ cINs from initial contact with a BV in brain slices from E13.5 GFP^+^cINs-control (open circles and triangles) or from GFP^+^cINs-OPC-depleted embryos (purple circles and triangles). Triangles correspond to brain slices treated with anti-Cxcl12 blocking antibody. Circles correspond to slices treated with vehicle (n = 24-25 cells from 3 embryos from 3 dams; two-way ANOVA; ***p=0.0006 after 90 min, ****p<0.0001 after 120 to 180 min). (G) Distribution of average migration speed of Dlx5,6-GFP^+^ cINs moving along cortical BVs on Cxcl12-treated (open and purple triangles) or vehicle-treated (open and purple circles) organotypic brain slices from E13.5 GFP^+^cINs-control and GFP^+^cINs-OPC-depleted embryos (n = 25-26 cells from 3 embryos, from at least 3 dams; one-way ANOVA; ****p<0.0001). (H) Histogram representing the displacement of Dlx5,6-GFP^+^ cINs after initial contact with a BV at time 0, on brain slices from E13.5 GFP^+^cINs-control vehicle-treated (open circles) and GFP^+^cINs-control XAV939-treated embryos (red circles) (n = 23-28 cells from 3 embryos, from at least 3 dams; two-way ANOVA; ****p<0.0001 after 30 to 180 min). (I) Distribution of average migration speed of Dlx5,6-GFP^+^ cINs moving along cortical BVs on organotypic brain slices from E13.5 GFP^+^cINs-control vehicle-treated (open circles) or GFP^+^cINs-control XAV939-treated embryos (red circles) (n = 24-28 cells from 3 embryos, from at least 3 dams; unpaired t-test; ****p<0.0001).

### vOPCs modulate cINs migration in the forebrain by unidirectional contact-repulsion

Loss of vOPCs affects both forebrain distribution and migration of contemporary cINs, thereby suggesting that these cells may crosstalk to control their migration. To address this question, we co-cultured MGE explants from E13.5 Nkx2.1:Cre; R26^CAGTdTomato^ embryos almost exclusively populated by TdTomato-positive (Tom^+^) cINs in compartmented dishes (that prevent physical interaction in a share culture medium) with homochronic WT cortical feeder or dissociated POA isolated from E13.5 SOX10:GFP-DTA embryos and containing only GFP^+^ vOPCs (Figures 5A–5D). The number and dispersion of migrating Tom^+^ cINs around their MGE core explant were comparable in both co-culture conditions, suggesting that vOPCs repel cINs via cell-cell interaction rather than by releasing chemorepulsive cues (Figures 5E–5H). In order to test this hypothesis, we analysed the migration between Tom^+^cINs moving away from Nkx2.1:Cre; R26^CAGTdTomato^ MGE explants with GFP^+^ vOPCs exiting Sox10:GFP-DTA POA explants on homochronic cortical feeder (Figure 5I). The number of migrating Tom^+^cINs invading Sox10:GFP-DTA POA explants was reduced after 48h of culture, as compared to those isolated from vOPC-depleted embryos. This suggested that vOPCs-enriched POA prevents cINs invasion (number of cINs in POA control explants: 2.5 ± 1.5 and number of cINs in OPC-depleted POA explants: 20.75 ± 5.36, p=0.029; Figures 5J–5L). At cellular level, super-resolution live imaging showed that vOPCs-cINs contact is a two-step event during which vOPCs extend one branch that recognises cINs leading process on its growth cone-like structure followed by polarity reversal of cINs (Figure 5M; Supplemental movie S4). This leads to a dramatic change in the migration trajectory of cINs without reciprocity and alteration of vOPCs motion (Figure 5M). This unidirectional contact-repulsion (UCoRe; from vOPCs to cINs) occurred in average 78.6min ± 6.11min after the initial contact (Figure 5N) and is highly specific as it only occurs between vOPCs and cINs (97.1% of the contacts with 71.4 % leading to complete cINs polarity reversal, the remaining ones are angled repulsion) (Figure 5O) and also occurs in organotypic brain slice, a close to physiological context (Supplemental movie S5). We also observed self-repulsion between vOPC pairs (39.0% of the recorded contacts) or cIN pairs (8.7% of the recorded contacts) (Figures 5P, 5Q).

**Figure 5.**
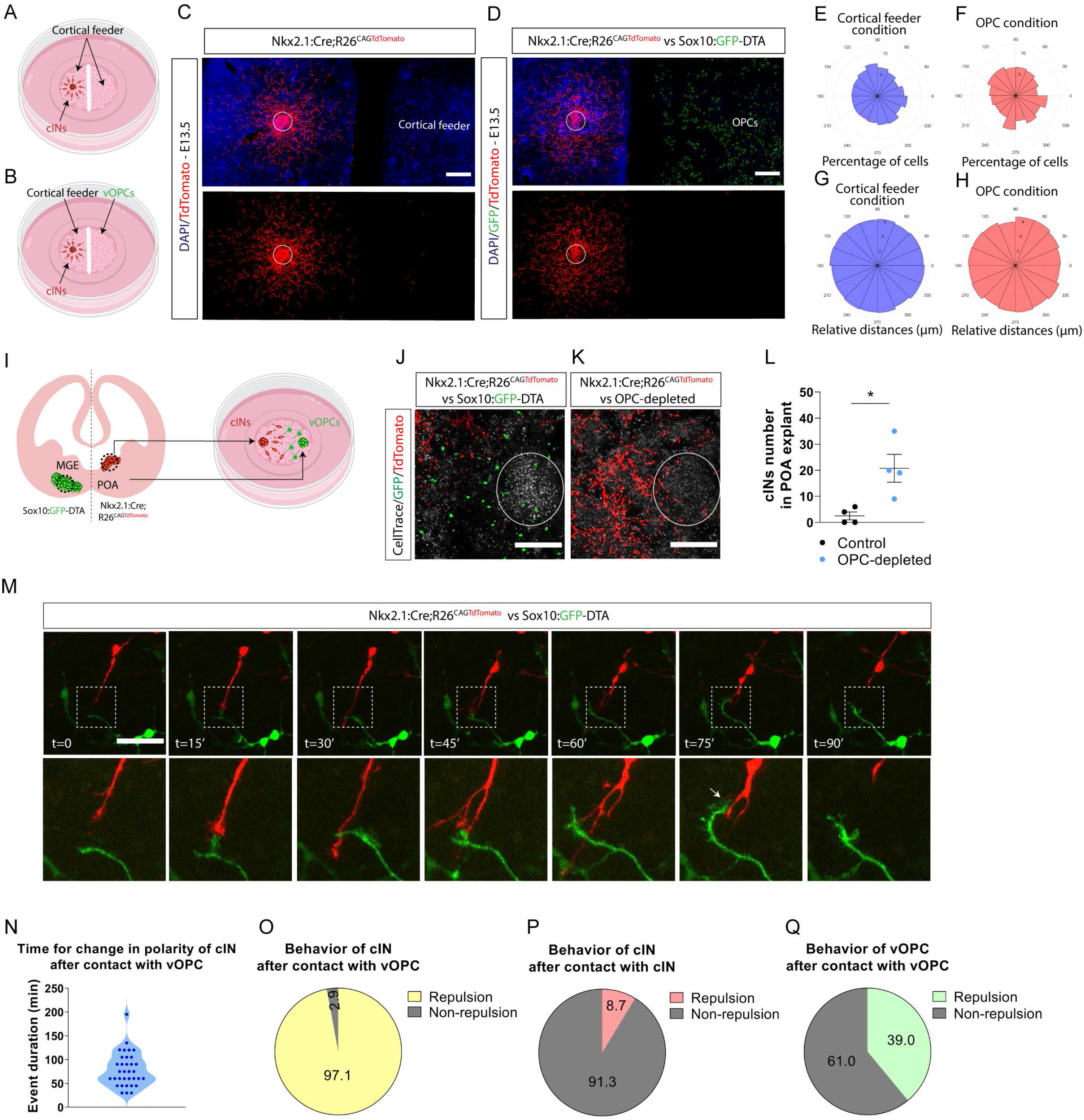
Analysis of vOPCs and cINs interaction. (A-B) Experimental culture set up to analyze vOPCs secretome influence on cINs migration. cINs explants from Nkx2.1:Cre;R26^CAGTdTomato^ embryos were cultured on mixed cortical feeder in one side of a bi-compartmented dish, while the other side was seeded with either mixed cortical feeder (A) or dissociated POA enriched in vOPCs and isolated from E13.5 Sox10:GFP-DTA embryos (B). (C-D) Representative images illustrating the co-cultures depicted in (A) and (B), respectively. Scale bar: 500μm. (E-H) Phase histograms showing the fraction (E, F) or the distance travelled (G, H) by cINs from their explant core in the culture set up depicted in (A-B). (I) Experimental *in vitro* set up to assess interaction between vOPCs (green, isolated from E13.5 Sox10:GFP-DTA POA) and cINs (red, isolated from E13.5 Nkx2.1:Cre;R26^CAGTdTomato^ MGE). (J-K) Representative images of co-cultures of MGE-derived explants isolated from organotypic brain slices of E13.5 Nkx2.1:Cre;R26^CAGTdTomato^ (enriched in TdTomato^+^ cINs) embryos facing POA-derived explants obtained from organotypic brain slices of E13.5 Sox10:GFP-DTA (J) or OPC-depleted (K) embryos. Scale bar: 200μm. (L) Absolut numbers of cINs invading POA-derived explant from Sox10:GFP-DTA (black circles) or OPC-depleted (blue circles) embryos (n = 4 explants from 3 embryos, from at least 3 dams; Mann-Whitney test; *p=0.0286). (M) Time-lapse video microscopy sequence illustrating the physical interaction between a TdTomato^+^ cIN (red), and a GFP^+^ vOPC (green). The white arrow depicted at t=75’ points the physical contact between the cIN and the vOPC. Scale bar: 200μm. (N) Violin plot indicating the time for change in behavior of cINs after contacting a vOPC (78.60 ±6.11). (O-Q) Pie charts indicating the frequency of repulsive and neutral behaviors of cINs after contacting vOPCs (O) or other cINs (P), or between vOPCs (Q); repulsive behaviors are represented in yellow (O), red (P) or green (Q), while neutral behaviors are represented in grey.

Altogether, these data suggest that vOPCs steer the forebrain migration of cINs via UCoRe, a peculiar mode of contact-repulsion. This influences the final distribution of cINs in the cortical parenchyma and allows vOPCs to establish a privileged interaction with BVs to which both cINs and vOPCs share common attraction as a result of released endothelial Cxcl12.

## Discussion

Here, we provide an in-depth characterization of the migration pattern of first-wave vOPCs in the developing mouse brain. Despite being born contemporaneously in shared subpallial germinal regions, vOPCs and cINs use distinct migration modes and occupy mutually exclusive forebrain territories while moving towards the cortex. Moreover, these cell types undergo saltatory migration characterized by different kinetics and, while vOPCs continuously screen blood vessels (BVs) to invade the forebrain, cINs navigate in organized streams. Cell migration is regulated by guidance cues such as Cxcl12, a chemokine to which both cINs and vOPCs are responsive. The release of Cxcl12 by intermediate progenitors in the cortex streams the migration of cINs within IZ/SVZ ^20, 22^. This chemokine is also secreted by forebrain BVs that attract vOPCs^13^. While genetic ablation of cINs does not affect the cortical distribution of vOPCs, loss of vOPCs impairs the migration of cINs in the forebrain, further suggesting that cINs require vOPCs to properly navigate to and within the developing cerebral cortex. We showed that vOPCs ablation increases the local availability of Cxcl12 in the vicinity of BVs to which cINs are now attracted, thereby altering their migration pattern. At the cellular level, we demonstrate that vOPCs impose unidirectional contact-repulsion (UCoRe) to cINs, thus acting as “herd dogs” to prevent their interaction with BVs and to contribute to their overall guidance into the forebrain and toward their final cortical location. This mode of contact-repulsion with polarity reversal is very specific to vOPC-cIN pairs and distinct from self-repulsion, as previously described for OPCs themselves ^23^ and Cajal Retzius cells ^24, 25^. We are currently investigating the molecular basis of UCoRe. The reduction of cINs number in the cortex upon depletion of first-wave vOPCs is still observed around birth, while second-wave OPCs are actively migrating. This unexpected observation suggests that second-wave vOPCs functionally differ from first-wave vOPCs and may not perform UCoRe. Interestingly, cINs depletion does not have a major impact in the generation, survival and migration of first and second vOPC waves. However, by releasing Sonic hedgehog, cINs control the generation of dorsally-generated OPCs ^26^. This is another example strengthening the concept that cINs-OPCs crosstalk varies in space and time. Importantly, vOPCs and cINs develop after birth specific interaction to form transient synapses and later engage into myelination of specific local circuits ^5, 11^. This suggests that: i/ UCoRe only occurs during development to specifically organize cIN migration; ii/ postnatal vOPCs and cINs express recognition molecular codes to promote longer term interaction.

Here we show that UCoRe is an efficient strategy to organize the migration of distinct cell populations that use a common chemoattractant. It is particularly important to avoid the rerouting of cINs towards BVs where they slow down and aggregate. UCoRe is likely to be of growing importance to guide cINs as cortex mature and gets more perfused by vessels. Despite being attracted to BVs by locally secreted Cxcl12, it remains unclear why the migration of cINs, but not vOPCs, is reduced upon interaction with the forebrain vasculature, which remained morphologically unaffected upon loss of vOPCs. It is noteworthy that both Cxcr7 and Cxcr4 are responsive to Cxcl12 and their respective activation is required for proper migration of cINs in the cortex ^27, 28^. This does not seem to be the case for vOPCs that express low amount of Cxcr7 as compared to Cxcr4 ^13, 29^, and whose activation is not triggering a motogenic response ^30^. Thus, one hypothesis may be that blockade of Cxcr7 in cINs by BV-released Cxcl11 ^31, 32^, would reduce its motogenic activity in response to ambient Cxcl12, thereby slowing down cINs motion.

Altogether, our results show that despite being less numerous, vOPCs shape the migration of cINs across cortical territories by using UCoRe that prevent the attraction of cINs to BV-secreted Cxcl12 and that help them to integrate cortical streams filled by a gradient of Cxcl12 released by intermediate progenitors and meningeal cells ^27, 33^.

## Figure Legends

**Figure S1.**
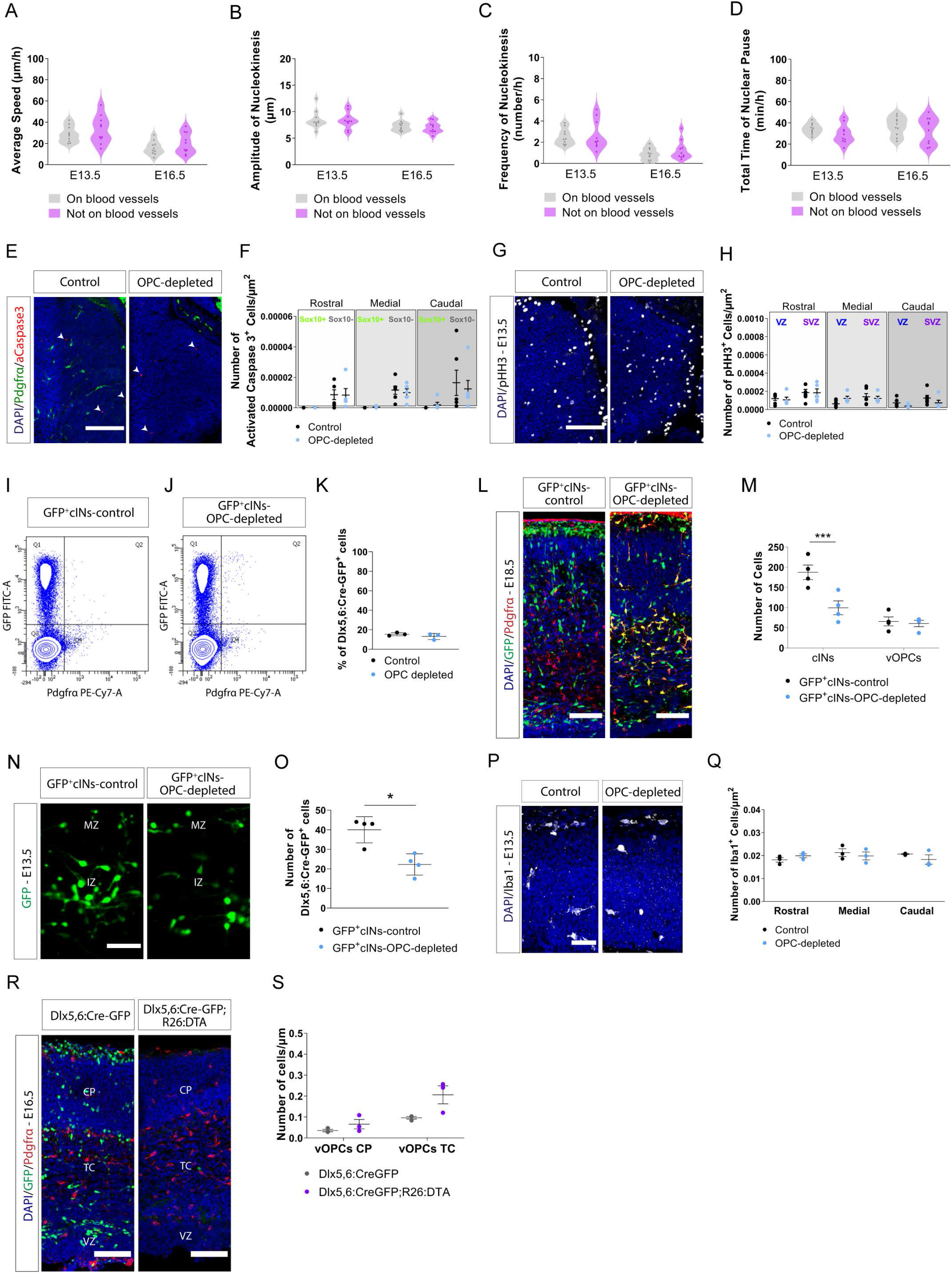
Migration properties of vOPCs and impact of vOPCs depletion on the generation and survival of cINs. (A-D) Distribution of average speed (A), amplitude of nucleokinesis (B), frequency of nucleokinesis (C) and total time of nuclear pause (D) of vOPCs migrating on BVs (gray) or not on BVs (purple) at E13.5 or E16.5 (n = 10 cells per group from 3 embryos, from at least 3 dams; two-way ANOVA). (E) Immunolabeling to detect expression of Pdgfrα (green) and activated caspase 3 (aCaspase3 - red). White arrowheads show dying aCaspase3^+^ cells in the subpallium of E13.5 Control (left) or OPC-depleted (right) embryos. Nuclei are counterstained with DAPI. Scale bar: 50μm. (F) Quantification of the number of aCaspase3^+^; Sox10^+^ cells and aCaspase3^+^; Sox10^-^ cells in rostral, medial and caudal sections of the subpallium of E13.5 Control (black circles) and OPC-depleted (blue circles) embryos (n = 5 to 6 embryos from at least 3 females; two-way ANOVA). (G) Immunolabelings to detect expression of pHH3 (white) in the subpallium of E13.5 Control (left) and OPC-depleted (right) embryos. Scale bar: 50μm. (H) Quantification of pHH3^+^ cells number in the VZ and SVZ of rostral, medial and caudal sections of the subpallium from E13.5 Control (black circles) and OPC-depleted (blue circles) embryos (n = 5 to 6 embryos from at least 3 females; two-way ANOVA). (I-J) FACS of Dlx5,6-GFP^+^ cINs dissociated from the brain of E13.5 GFP^+^cINs-control (I) or GFP^+^cINs-OPC-depleted embryos (J) identified in Q1 by high GFP expression and low Pdgfrα one. (K) Corresponding histogram comparing percentage of FACSed Dlx5,6-GFP^+^ cINs in GFP^+^cINs-control (black circles) or GFP^+^cINs-OPC-depleted (blue circles) (n = 3 embryos from 3 females; Mann-Whitney test). (L) Immunolabeling to detect expression of GFP (green) and Pdgfrα (red) in E18.5 GFP^+^cINs-control or GFP^+^cINs-OPC-depleted embryos. In GFP^+^cINs-control embryo, vOPCs are stained in red and cINs in green. In GFP^+^cINs-OPC-depleted embryo, vOPCs are yellow and cINs are green. The number of cINs is decreased upon first-wave vOPCs elimination. Nuclei are counterstained with DAPI. Scale bar: 100μm. (M) Quantification of the number of cINs and vOPCs in the cortex from E18.5 GFP^+^cINs-control (black dots) or GFP^+^cINs-OPC-depleted (blue dots) embryos (n = 4 embryos from 3 females; two-way ANOVA, ***p=0.0008). (N) Magnified area on organotypic brain slices from a E13.5 GFP^+^cINs-control (left) or GFP^+^cINs-OPC-depleted embryo (right) showing the distribution of Dlx5,6-GFP^+^ cINs. MZ and IZ correspond, respectively, to the migratory stream in the marginal zone and intermediate zone. Scale bar: 50μm. (O) Histogram representing the number of Dlx5,6-GFP^+^ cINs in brain slices from E13.5 Control (black circles) or OPC-depleted (blue circles) embryos (n = 4 embryos from at least 3 females; Mann-Whitney test; *p=0.0286). (P) Immunolabeling of Iba1^+^ microglia on brain slices from E13.5 Control or OPC-depleted embryos. Scale bar: 50μm. (Q) Quantification of the number of Iba1^+^ microglia in the cortex of E13.5 Control (black circles) and OPC-depleted (blue circles) embryos at rostral, medial and caudal levels (n = 3 embryos from 3 females; two-way ANOVA). (R) Immunolabeling to detect expression of GFP (green) and Pdgfrα (red) in E16.5 Dlx5,6:Cre-GFP or Dlx5,6:Cre-GFP;R26:DTA embryos. Nuclei are counterstained with DAPI. Scale bar: 100μm. (S) Quantification of the number of Pdgfrα^+^ cells normalized by cortical thickness in E16.5 Dlx5,6:CreGFP (grey circle) or Dlx5,6:CreGFP;R26:DTA (purple circle) embryos (n = 3 embryos from 3 females; two-way ANOVA).

**Figure S2.**
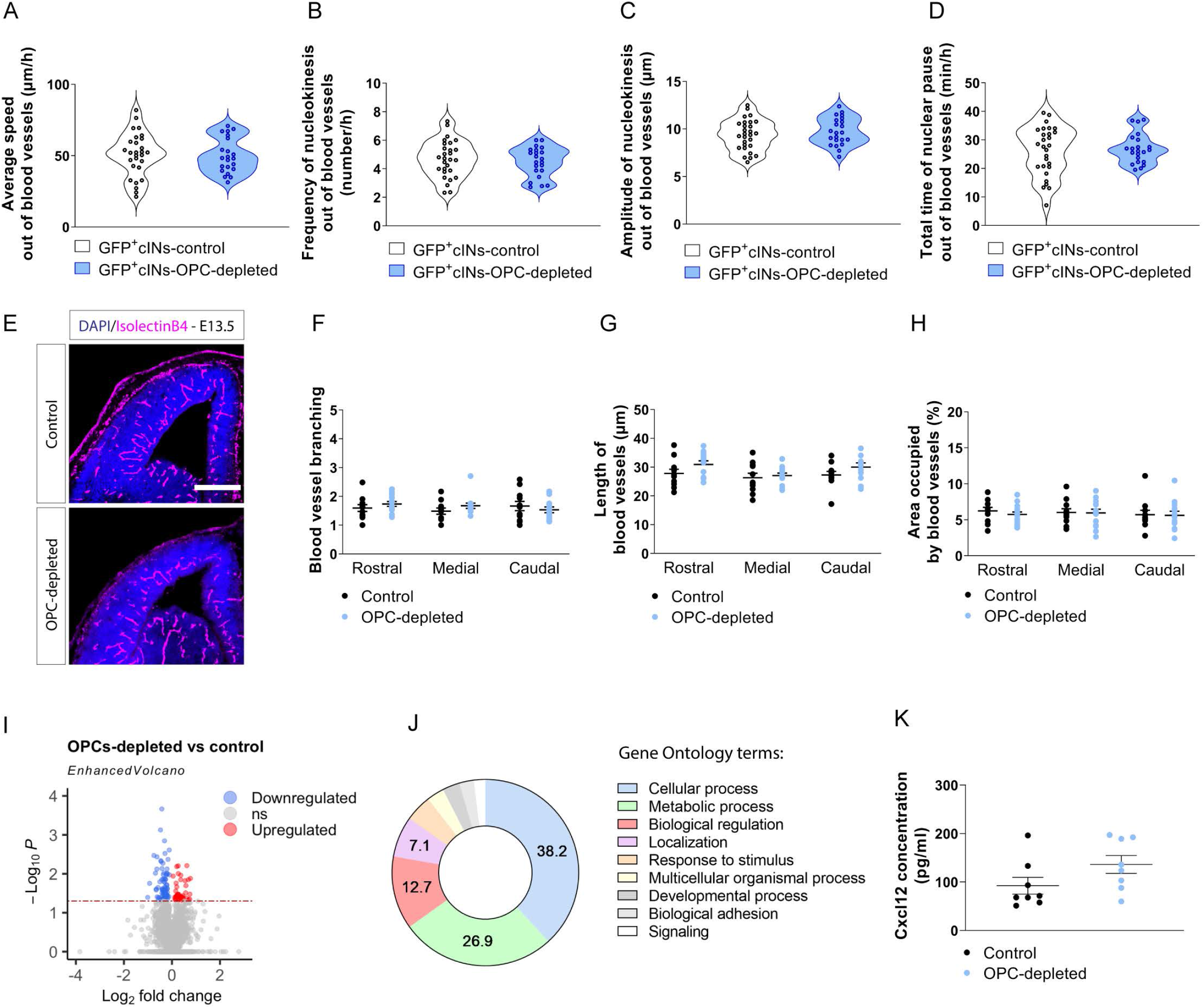
Migrating properties of cINs that are not moving along BVs, whose organization does not change upon depletion of vOPCs. (A-D) Distribution of average speed (A), frequency of nucleokinesis (B), amplitude of nucleokinesis (C) and total time of nuclear pause (D) of recorded Dlx5,6-GFP^+^ cINs migrating in cortical parenchyma region deprived of BVs in organotypic brain slices from E13.5 GFP^+^cINs-control or GFP^+^cINs-OPC-depleted embryos (n = 22 to 28 cells from at least 3 embryos from 3 dams; Unpaired t-test). (E) Coronal sections of a Control and OPC-depleted embryo stained with isolectinB4. Scale bar: 300μm. (F-H) Histograms representing branching (F), length (G) and cortical area covered (H) by BVs in the cortex of E13.5 Control (black circles) and OPC-depleted (blue circles) embryos in rostral, medial and caudal levels (n = 11 to 13 embryos from at least 3 females; two-way ANOVA). (I) Volcano plot indicating the significantly upregulated (red) and downregulated (blue) proteins identified by label-free differential proteomics (p-value cut-off <0.05; Welch t-test). (J) Gene ontology term identification and enrichment analysis. (K) Histogram depicting the quantification of Cxcl12 measured by ELISA in the cortex of E13.5 Control (black circles) and OPC-depleted (blue circles) embryos (n = 8 embryos from 4 dams; Mann-Whitney test).

***Movie S1. Migration of vOPCs in the MGE/POA territory they occupy at E13.5, Related to Figure 1*.** Time lapse video microscopy performed on organotypic slices from E13.5 Sox10:GFP-DTA mice showing the migration of GFP^+^ vOPCs in MGE/POA territory. VZ: ventricular zone, SVZ: subventricular zone, MZ: mantle zone. Duration of recording is 10h.

***Movie S2. Migration of cINs in the developing cortical wall at E13.5, Related to Figure 2A*.** Time lapse video microscopy performed on organotypic slices from E13.5 Dlx5,6:Cre-GFP mice showing the migration of cINs in the developing cortical wall. cINs migrate within their two main tangential streams, the MZ and IZ. Duration of recording is 5h.

***Movie S3. Migration of vOPCs in the developing cortical wall at E13.5, Related to Figure 2A*.** Time lapse video microscopy performed on organotypic slices from E13.5 Sox10:GFP-DTA mice showing the migration of vOPCs in the developing cortical wall. vOPCs do not display stream organization. Duration of recording is 5h.

***Movie S4. UCoRe between E13.5 cIN and vOPC on homochronic mixed cortical feeder, Related to Figure 5*.** Time lapse video microscopy showing repulsion between cIN and vOPC on homochronic explant culture. The arrow shows the contact between the growth cones of cIN and vOPC.

***Movie S5. UCoRe between E13.5 cIN and vOPC on organotypic brain slice, Related to Figure 5*.** Time lapse video microscopy showing repulsion cIN (Nkx2.1:Cre; R26^CAGTdTomato^) and vOPC (Sox10:Venus) on organotypic slices. The arrow shows the contact between the growth cones of cIN and vOPC. BVs are stained with isolectinB4 (white).

## Material & Methods

### KEY RESOURCES TABLE

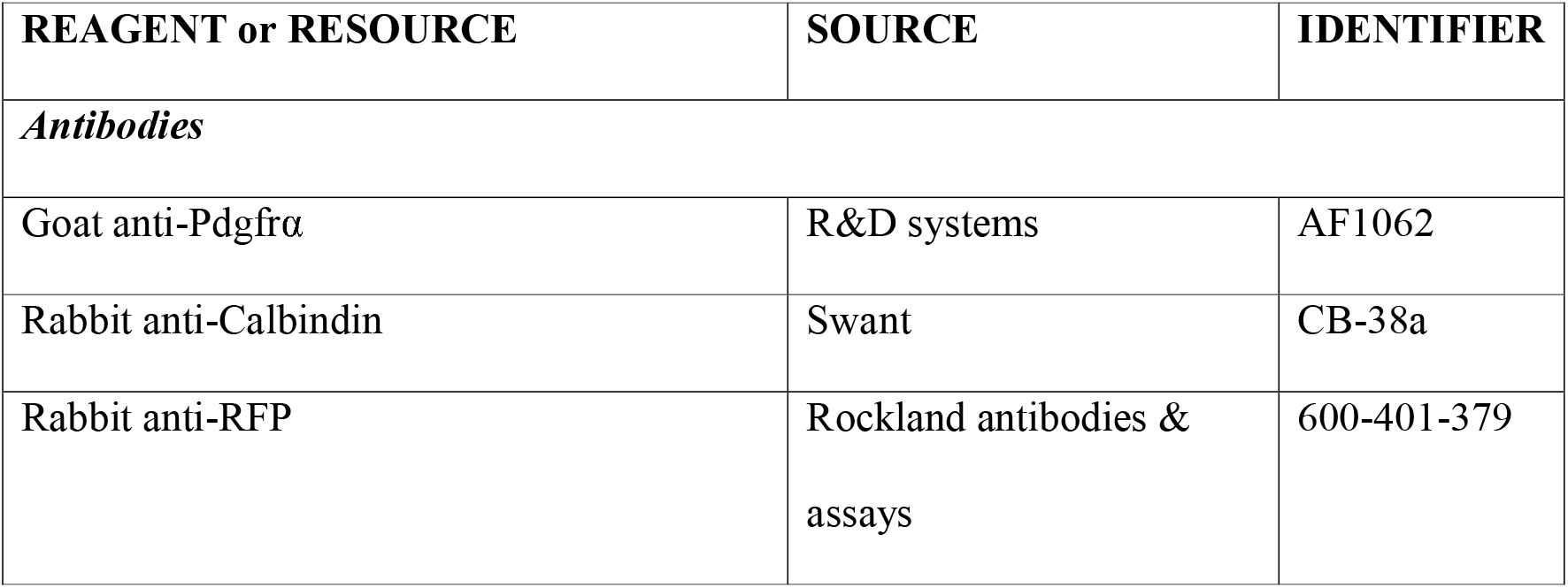
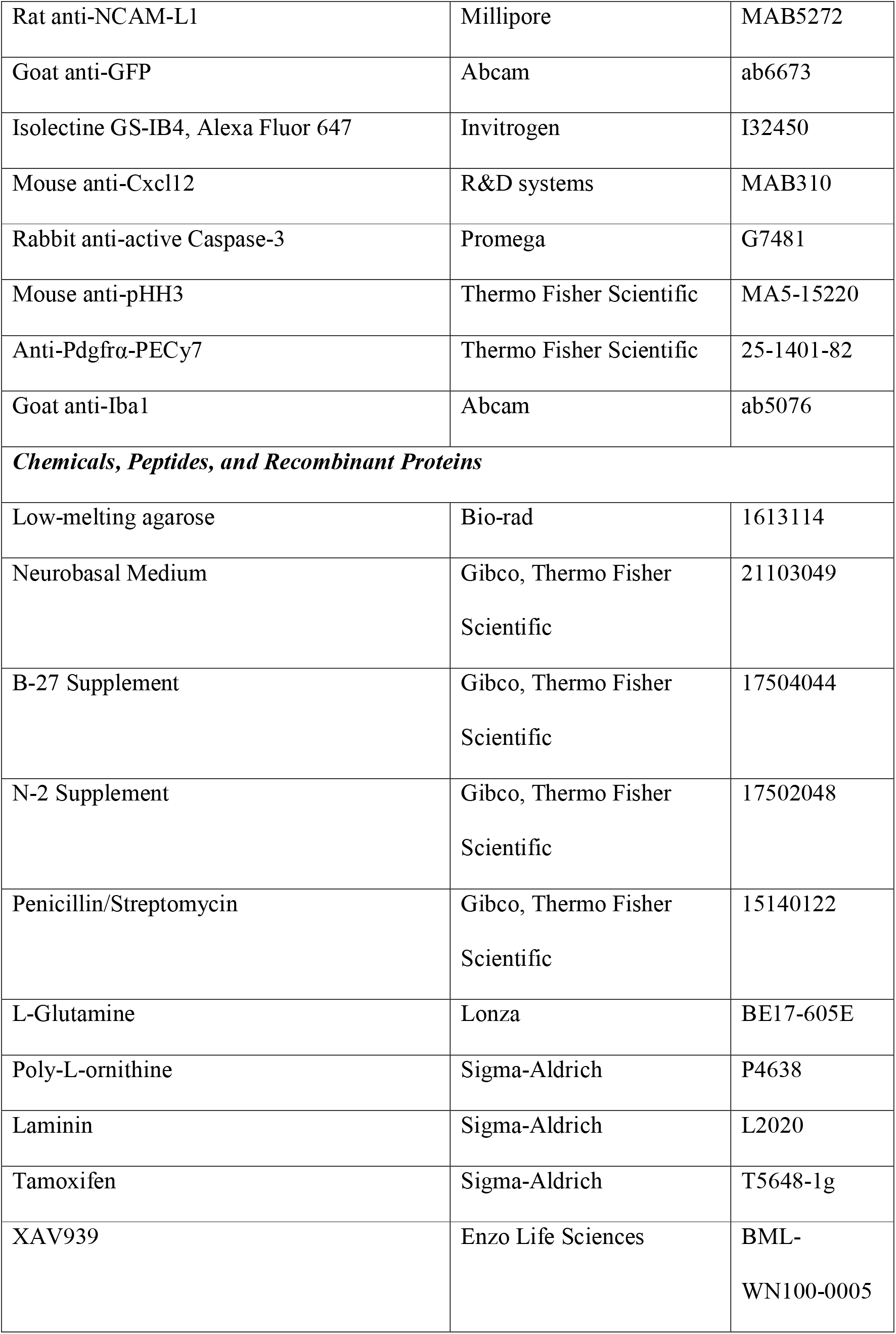
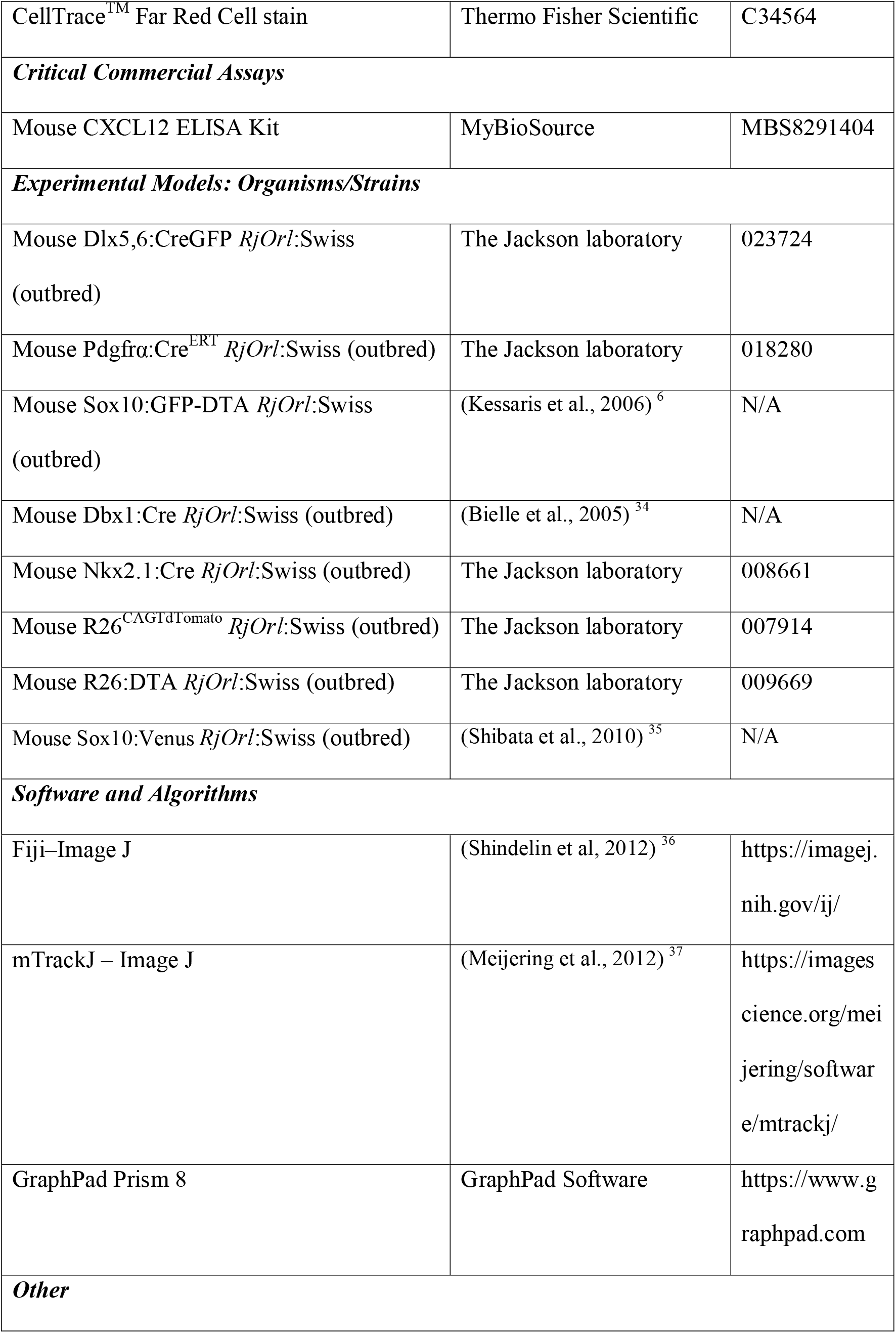
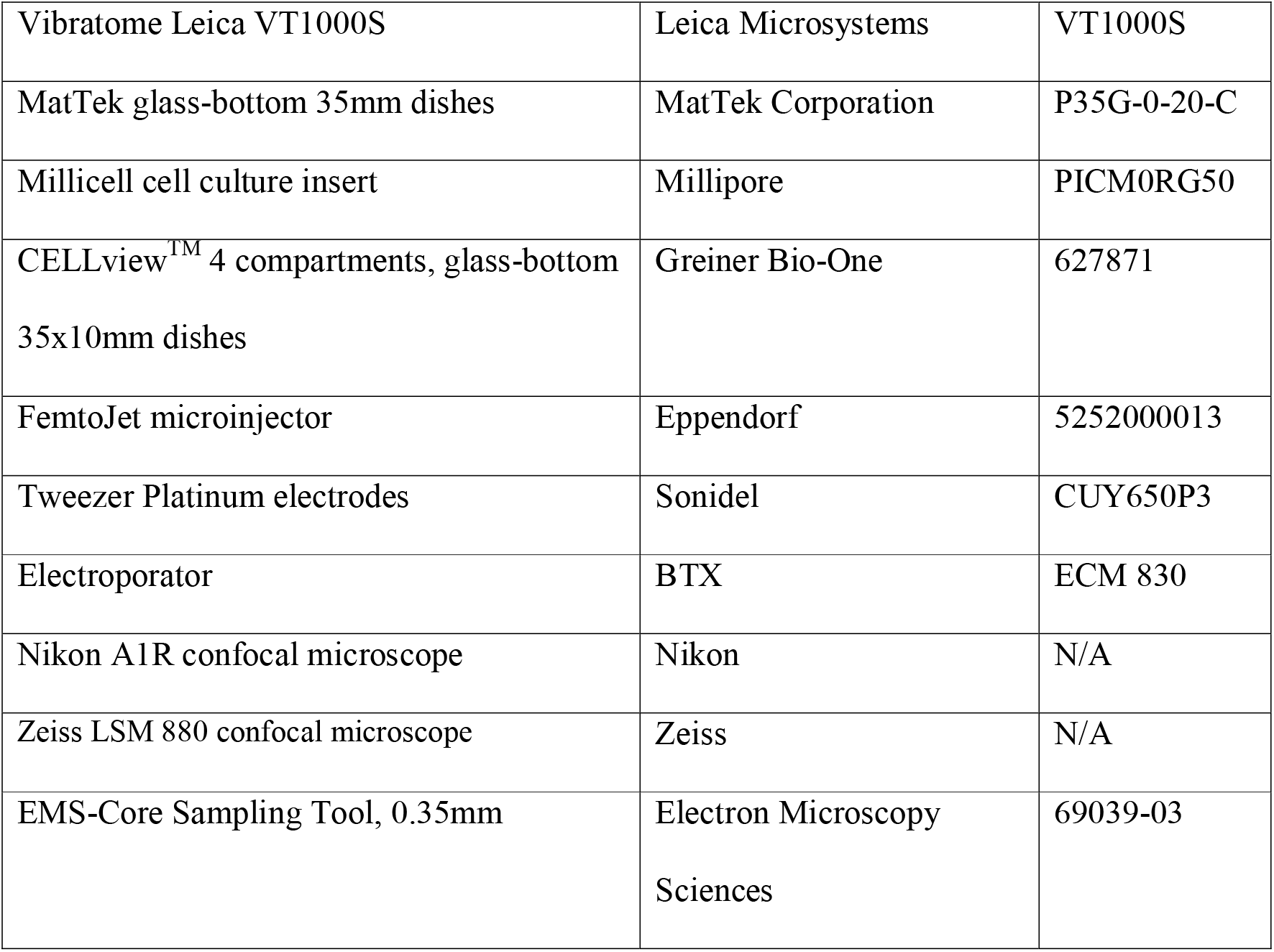

### CONTACT FOR REAGENT AND RESOURCE SHARING

Further information and requests for reagents may be directed to and will be fulfilled by the Lead Contact, Laurent Nguyen.

### EXPERIMENTAL MODEL AND SUBJECT DETAILS

#### Mouse genetics

All mice were handled according to the ethical guidelines of the Belgian Ministry of Agriculture in agreement with European Community Laboratory Animal Care and Use Regulations (86/609/CEE, Journal Officiel des Communautées Européennes, L358, 18 December 1986) and with our experimental protocol n°17-1966. All mouse models were maintained in SWISS genetic background. Both males and females were used for all analyses. Wild-type SWISS pregnant females were purchased from Janvier-Labs (France) for the analysis of cINs and vOPCs distribution in the forebrain from E11.5, E13.5, and E16.5 embryos. Dbx1:Cre (kindly provided by Dr. Alessandra Pierani) or Nkx2.1:Cre (The Jackson Laboratory, stock number 008661) mice were time-mated to R26CAGTdTomato (The Jackson Laboratory, stock number 007914) mice. E13.5 embryos were used to visualize recombined cells and to analyze vOPCs progeny. Sox10:GFP-DTA (kindly provided by Dr. Emmanuelle Huillard), Sox10:Venus (kindly provided by Dr. Hideyuki Okano) as well as Dlx5,6:CreGFP (The Jackson Laboratory, stock number 023724) transgenic models were used at E13.5 and E16.5 to characterize the migration of vOPCs or cINs respectively. Sox10:GFP-DTA females were also crossed with either Pdgfra:Cre^ERT^ (The Jackson Laboratory, stock number 018280) or Dlx5,6:CreGFP;Pdgfra:Cre^ERT^ males to deplete first-wave vOPCs by tamoxifen injections at E11.5 and E12.5. Finally, cINs depletion was performed by mating R26:DTA females (The Jackson Laboratory, stock number 009669) to Dlx5,6:CreGFP males.

### METHODS DETAILS

#### Organotypic slice culture from embryos brains and time-lapse recordings

For a detailed protocol of organotypic brain slice culture experiment, please refer to ^38^. Briefly, heads from E13.5 or E16.5 embryos were harvested in ice-cold HBSS containing Ca^2+^ and Mg^2+^ and embedded in 4% low-melting agarose diluted in HBSS. Brains were cut into 300μm coronal slices using a Leica VT1000S vibratome set with the following parameters: section thickness: 300 μm; speed: 3; frequency: 3. Slices were immediately harvested on 0.4μm millicell cell culture insert placed on MatTek dishes filled with supplemented Neurobasal medium (NBM). To stain BVs, 2μl of isolectinB4 was added per slice. Slices were kept in a humidified incubator saturated with 5% CO_2_ for maximum 8h before time-lapse recording.

For the analysis of vOPCs and cINs migration, recordings were performed for at least 5h using a Nikon A1R inverted confocal microscope. Acquisitions were obtained each 5 min with an average of 5 z-steps. For Cxcl12 blockade, 5μL of Cxcl12 antibody diluted in DPBS (100μg/mL) was added on the top of organotypic slices for 2h before time-lapse recording. For Wnt signaling inhibition, 5μL of XAV939 diluted in NBM (100nM) was added on the top of organotypic slices for 8h before time-lapse recording. XAV939 was also added at the same concentration to the culture medium. Control organotypic slices were treated with Cxcl12 or XAV939 vehicle diluted in NBM (DPBS or DMSO, respectively). For all the experiments, the dishes were maintained at 37°C in a humidified chamber saturated with 5% CO2.

#### MGE and POA explant culture on homochronic cortical feeder for time-lapse recordings

CELLviewTM compartmented glass-bottom dishes were coated with poly-ornithine (0.1 mg/mL) for 45 min at 37°C, followed by an incubation with laminin (5 μg/mL) for 2h at 37°C. A feeder layer of cortical cells was prepared by mechanical dissociation of E13.5 wild-type cortices (2 embryos/dish) using P1000 pipette. The dissociated cells were plated in coated glass-bottom dishes and were left adhere for 2h at 37°C in a humidified incubator saturated with 5% CO2. NBM supplemented with B27 containing vitamin A (2%), penicillin/streptomycin (1%) and L-glutamine (1%) was slowly added to the cells, and the dishes were kept in the incubator for 24h. The MGE explants from Nkx2.1:Cre;R26^CAGTdTomato^ embryos were then plated on the top of the cortical feeder layer together with the POA explants from Sox10:GFP-DTA embryos. Brains were dissected in ice-cold HBSS containing Ca^2+^ and Mg^2+^ and embedded in 4% low-melting agarose diluted in HBSS. Brains were cut into 300μm coronal slices using a Leica VT1000S vibratome set with the following parameters: section thickness: 300 μm; speed: 3; frequency: 3. Medial to caudal slices were placed in ice-cold HBSS in a Petri-dish coated with a thin layer of low-melting agarose. MGE and POA explants were obtained using a 350μm punch, then plated on the top of the cortical feeder layer. For the analysis of contact/repulsion between cINs and vOPCs, recordings were performed for at least 10h using a Nikon A1R or a Zeiss LSM 880 inverted confocal microscope. Acquisitions were obtained each 5 min with an average of 15 z-steps. For all the experiments, the dishes were maintained at 37°C in a humidified chamber saturated with 5% CO2.

#### Brain tissue collection and preparation

After dissection, embryonic heads or brains were fixed in 4% paraformaldehyde (PFA) overnight (ON) at 4°C. The brains were dehydrated in a 20% sucrose solution diluted in PBS ON at 4°C. After complete dehydration, brains were embedded and froze in OCT using dry ice. Brains were kept at −80°C until their cutting at the cryostat (20μm or 80μm slices).

#### Immunohistochemistry on brain slices

Brain slices were permeabilized and blocked in PBS containing 0.3% Triton X-100 and 10% donkey serum. Slices were then incubated ON at 4°C with primary antibodies diluted in PBS containing 0.3% Triton X-100 and 1% donkey serum. Primary antibodies were used at the following concentrations: goat anti-Pdgfrα 1:500, rabbit anti-Calbindin 1:1000, rabbit anti-RFP 1:1000, rat anti-NCAM-L1 1:1000, goat anti-GFP 1:1000, Isolectine GS-IB4 1:1000, rabbit anti-active Caspase-3 1:500, mouse anti-pHH3 1:500, goat anti-Iba1 1:1000. Images were acquired using a Nikon A1R inverted confocal microscope with a 10x or 20x objective.

#### Immunocytochemistry on MGE explants

Cells were fixed in 4% PFA for 20 min, and permeabilized in PBS containing 0.3% Triton X-100 and 10% donkey serum for 1h at room temperature. Cells were then incubated ON at 4°C with primary antibodies diluted in PBS containing 0.3% Triton X-100 and 1% donkey serum. Primary antibodies were used at the following concentrations: rabbit anti-RFP 1:1000, goat anti-GFP 1:1000. Images were acquired using a Nikon A1R inverted confocal microscope with a 10x objective.

#### Fluorescent Activated Cell Sorting

E13.5 whole forebrains were dissociated with papain containing 50mM DNase I. Cells were filtrated through a 40μm filter, and stained with anti-PDGFRα-PECy7 antibody (1:20) for 1h at 4°C before the phenotyping analysis. Percentage of GFP+/Pdgfrα-cells was calculated using FACS ARIA III with FACSDiva 8.0 software (125.000 events).

#### Elisa assay

Cxcl12 levels in dissected E13.5 cortices were quantified following manufacturer’s recommendations.

#### Sample preparation for proteomic analysis

Cortices from E13.5 Control and OPC-depleted embryos were dissected and froze using liquid nitrogen. The samples were solubilized in Tris HCl 10 mM pH 7.4, SDS 4%, Protease Inhibitor Cocktail EDTA-free (Complete, Sigma Aldrich). 5 units of DNAse were added by 100 mg of tissue to each sample. The samples were submitted sonication for 3 times 20 seconds, then let 10 minutes under agitation at 4°C. The protein content of the samples was then quantified using RCDC kit (Biorad). For each sample, 20 μg of proteins were diluted in 50 mM final concentration of ammonium bicarbonate at a protein concentration of 0.75 μg/mL then reduced using dithiothreitol (DTT), alkylated with iodoacetamide. Then the 2D-Clean up kit (GE) was applied according to manufacturer recommendations, to eliminate impurities not compatible with mass spectrometry analysis. The protein pellets after the washing steps were further resolubilized in bicarbonate ammonium 50mM. The samples were digested in solution with trypsin (16 hours at 37°C ratio tryspin/total proteins (W/W) (1/50), 3h at 37°C with ratio 1/100 in 80% ACN). The reaction was stopped by addition of trifluoroacetic acid acid. The samples were evaporated to dryness in a speed vacuum. For each sample, a quantity of 3.5 μg of protein digest was purified on a Ziptip C18 (Millipore), dried and re-suspended in 100 mM Ammonium formiate (pH 10) at 0.333 μg/μL. Each sample was spiked with a commercial mixture of protein digest standards originated from non-human biological material: the MassPREP™ Digestion Standards (Waters, Corp., Milford, USA), at 150 fmol of ADH per sample. Sample injection order on 2D-nanoUPLC-ESI-Q orbitrap was randomized and 9 μL, corresponding to 2 μg were injected per sample. Samples were injected onto the 2D-nanoAquity UPLC (Waters, Corp., Milford, USA) coupled online with a Q Exactive Plus system (Thermo Scientific, USA) in nanoelectrospray positive ion mode. The configuration of the 2D-nanoUPLC system was a reversed phase pH 10 - reversed phase pH 3 based two-dimension separation. The first-dimension separation was made on an X-Bridge BEH C18 5 μm column (300 μm x 50 mm). The trap column Symmetry C18 5μm (180 μm x 20 mm) and analytical column HSST3 C18 1.7 μm (75 μm x 250 mm) (Waters, Corp., Milford, USA) were used after an online dilution to lower pH values. The samples were loaded at 2 μL/min (20 mM ammonium formate solution adjusted to pH 10) on the first column and subsequently eluted in three steps (13.3%, 19% and 65% acetonitrile) to the low pH columns. Each eluted fraction was desalted on the trap column after a ten times online dilution to pH 3 and subsequently separated on the analytical column; flow rate 250 nL/min, solvent A (0.1% formic acid in water) and solvent B (0.1% formic acid in acetonitrile), linear gradient 0 min, 99% A; 5 min, 93% A; 140 min, 65% A. The total run time of each step was 180 min. The mass spectrometer method is a TopN-MSMS method where N was set to 12, meaning that the spectrometer acquires one Full MS spectrum, selects the 12 most intense peaks in this spectrum (singly charged precursors excluded) and makes a Full MS2 spectrum of each of these 12 compounds. The parameters for MS spectrum acquisition are: Mass range from 400 to 1750 m/z, resolution of 70000, AGC target of 1e6 or maximum injection time of 200 ms. The parameters for MS2 spectrum acquisition are: isolation window of 2.0 m/z, normalized collision energy (NCE) of 25, resolution of 17500, AGC target of 1e5 or maximum injection time of 50 ms, underfill ratio of 1.0%.

#### Proteome analysis

For label-free quantification (LFQ) application, MaxQuant ver. 1.6.6.0 was used for the analysis of raw files. MS/MS spectra were analyzed using the Andromeda search engine and the following settings: database Uniprot reviewed Mus Musculus database (downloaded on the 23 March 2019, 17022 protein sequences) for interrogation, oxidation of methionines was set as variable modifications, carbamidomethylation of the cysteines as fixed modification. The maximum number of missed cleavages was set at two and the minimal peptide length for identification was set at 7 amino acids and at least two peptides per protein, including at least one unique peptide, were required for identification. Data normalization was performed using the LFQ algorithm^39^. The minimum ratio count for LFQ was set at 2. The main search tolerance was set at 4.5 ppm. Peptide spectrum match (PSM) and protein false discovery rates (FDR) were both set at 0.01. Data detected in at least 2 of the 3 samples per group were treated using Perseus ver. 1.6.6.0. Differentially expressed proteins were detected using a Welch t-test.

#### Time-lapse recording analysis

Average speed, amplitude and frequency of nucleokinesis, and total time of nuclear pause were analyzed using mTrackJ plugin on ImageJ software as reported in ^38^. The distances between cINs and blood vessels were manually measured using ImageJ software.

#### Histological analysis

Analysis of fluorescence intensity and quantification of cell numbers were performed by ImageJ software. Counting areas were adapted for each slice to fit the thickness of the cortex, with a fixed width of 300 μm at E13.5 and 400 μm at E16.5. Distances between VZ and IMS in E14.5 brains were manually measured using ImageJ software. Measurements of blood vessels branching and length were performed using Skeleton plugin in ImageJ software.

#### Quantification and statistical analyses

They were performed using GraphPad Prism 8. Statistical differences between two experimental groups were evaluated using parametric or non-parametric t-tests, after verifying the normality of distribution for each group. Multiple comparisons were performed using one way-ANOVA or two-way ANOVA. Results are expressed as mean ±SEM. The experimental n, statistical test used and statistical significance for each experiment are indicated in the figure legends.

## Author contributions

F.L., C.G., and L.N. designed the study. F.L. and C.G. performed and interpreted most experiments with L.N. G.W. and D.B. generate the proteomic data and analysed them. F.L., C.G., and L.N. wrote the manuscript with input from all co-authors.

### Competing interests

The authors declare that they have no competing interests.

## Acknowledgements

We thank the following colleagues for kindly providing transgenic mouse lines: Dr. Emmanuelle Huillard (Sox10:GFP-DTA), Dr. Hideyuki Okano (Sox10:Venus), Dr. Rachelle Franzen (Pdgfra:Cre^ERT^), Dr Alessandra Pierani (Dbx1:Cre). We thank Drs Maria Cecilia Angulo and Alessandra Pierani for providing a feedback on the manuscript as well as athe members of the Nguyen lab for their input. We thank Sandra Ormenese, Alexandre Hego, Céline Vanwinge, and Raafat Stephan from the GIGA-Cell Imaging and flow cytometry platforms as well as Gabriel Mazzucchelli and Dominque Baiwir from the GIGA-proteomics platform for their technical support.

F.L. and L.N. are respectively PhD student and Senior Research Associates from F.R.S-F.N.R.S. C.S. is postdoctoral fellow supported by the Belgian Science Policy (IAP-VII network P7/20. This work was made possible thanks to grant from the F.R.S.-F.N.R.S. (Synet; EOS 0019118F-RG36), the Fonds Leon Fredericq, the Fondation Médicale Reine Elisabeth, the Fondation Simone et Pierre Clerdent, the Belgian Science Policy (IAP-VII network P7/20), and the ERANET Neuron STEM-MCD and NeuroTalk.

## Notes

### Competing Interest Statement

The authors have declared no competing interest.

